# Mechanical control of the splicing factor PTBP1 regulates extracellular matrix stiffness-induced cell proliferation and mechanomemory

**DOI:** 10.1101/2024.05.07.592669

**Authors:** Pei-Li Tseng, Weiwei Sun, Ahmed Salem, Sarah Macfarlane, Annica K. B. Gad, Mark O. Collins, Kai S. Erdmann

## Abstract

Cells sense and respond to mechanical cues from their environment. Mechanical cues are important for many biological processes, including embryonic development, ageing, cellular homeostasis, and diseases. Cells translate mechanical cues into cellular biochemical signals that govern cellular behaviour, like cell proliferation or migration, via a process called mechanotransduction. However, this process and the proteins involved remain incompletely understood. Here, we present an unbiased and large-scale approach to identify proteins involved in mechanotransduction. The screen revealed that the splicing factor PTBP1 is a novel mechanotransducer. We show that the nuclear localisation of PTBP1 depends on extracellular matrix stiffness, cell density, and the actomyosin-based contractility of the cell. Furthermore, we demonstrate that PTBP1 promotes the mechanosensitive splicing of the adapter protein Numb and that alternative splicing of Numb is crucial for matrix stiffness-induced cell proliferation and mechanomemory. Our results support the idea that changes in alternative splicing are an integral part of mechanotransduction and provide a mechanism by which matrix stiffness regulates cell proliferation and the formation of a mechanomemory in cells.

## INTRODUCTION

Mechanical properties of the cellular microenvironment (e.g., the stiffness of the extracellular matrix (ECM), fluid flow, and pressure), play important roles in cell differentiation, homeostasis, embryonic development, as well as in disease progression such as cancer or fibrosis [1–3]. How mechanical cues are translated into biochemical cellular changes is still not fully understood. Plasma membrane resident mechanosensors such as integrins can sense extra- and intracellular forces and such mechanical forces are conducted through the cytoskeleton and the Linker of Nucleoskeleton and Cytoskeleton complex (LINC) to the nucleus leading to changes in chromatin organisation and nuclear pore complex permeability making the nucleus a central player in mechanotransduction [4–6]. Changes in chromatin structure can then have a direct effect on transcriptional activity whereas regulation of nuclear pore complex permeability allows shuttling of proteins between the nucleus and the cytoplasm depending on their molecular size and their mechanical stability [4, 7, 8]. Mechanoresponsive nuclear transport receptors like importin-7 have further been identified to drive nuclear import of transcriptional regulators that play an important role in mechanotransduction [9]. Thus, changes in the subcellular localisation of proteins between the cytosol and the nucleus is a fundamental principle of mechanotransduction [10, 11]. Proteins that undergo mechanosensitive shuttling between the cytosol and the nucleus include transcriptional regulators, such as the Yes 1 associated transcriptional regulator (YAP), the Myocardin related transcription factor A (MRTF-A), the Runt-related transcription factor 2 (Runx2), Snail and Twist [10–15]. Emerging evidence suggests that in addition to transcription factors also proteins that participate in other functions in the nucleus such as DNA-methylation can be regulated by mechanosensitive nuclear-cytoplasmic shuttling [16]. Furthermore, the Src homology region 2-domain containing protein tyrosine phosphatase 1 (SHP1) has been shown to undergo nuclear shuttling by directly forming a complex with YAP, which is important for controlling tyrosine phosphorylation events in the nucleus [17]. In order to identify proteins, which demonstrate a mechanosensitive regulation of their nuclear localisation we developed an unbiased screen combining the controlled regulation of cellular actomyosin contractility with *in situ* proximity biotinylation of the nuclear proteome, followed by mass spectrometry analysis [18]. Thereby, we identified the splicing regulator Polypyrimidine Tract Binding Protein 1 (PTBP1) as a novel mechanotransducer, whose subcellular localisation is mechanosensitive [19, 20]. We further show that PTBP1 regulates the alternative splicing of the adapter protein Numb in a mechanosensitive manner, which plays a critical role in ECM-stiffness induced cell proliferation. Furthermore, we observed that this mechanosensitive splicing event resulted in the formation of a mechanomemory with respect to cell proliferation on stiff matrix. These results support an emerging role of the regulation of alternative splicing of mRNAs as an integral part of mechanotransduction.

## RESULTS

### Design and validation of a screen for proteins with mechanosensitive nuclear localisation

To identify proteins with a potential role in mechanotransduction, we developed a screening approach based on the quantification of changes of the nuclear proteome upon acute changes in cellular actomyosin contractility as outlined in figure 1A, B. To increase the contractile force in cells we made use of the fact that activation of the small Rho GTPase RhoA results in myosin light chain phosphorylation and increased myosin activity, and therefore increased cytoskeletal actomyosin-based contractility [21, 22]. This has been exploited previously to increase cellular traction and intracellular tension of cells [23, 24]. We therefore created HEK293 cells that express a constitutive active variant of RhoA, under a tetracycline-inducible promoter (HEK293-tet-RhoA). We observed expression of this active variant of the RhoA protein, as early as 1 hour after tetracycline addition (Suppl. Fig. 1A, B). We further observed an increase in cellular traction after addition of tetracycline by using traction force microscopy (Suppl. Fig. 1C, D). To identify proteins enriched in the nucleus after the expression of the constitutive active RhoA expression we compared the nuclear proteome between cells which had been treated with tetracycline or mock treated for 2 hours. To avoid any changes to the nuclear proteome due to extensive mechanically handling of the cells using standard subcellular fraction methods we decided to adopt a proximity-dependent labelling approach to biotinylate nuclear proteins *in situ.* For this we stably transfected a recently engineered biotin-ligase (NL-TurboID) with increased enzymatic activity and harbouring a nuclear localisation signal [18] into HEK293-tet-RhoA cells (generating HEK293-tet-RhoA-TurboID). As expected, NL-TurboID was localised to the nucleus (Suppl. Fig. 1E). We established efficient biotinylation of proteins via western-blotting using two independent HEK293-tet-RhoA-TurboID clones (Suppl. Fig. 2A). The transcriptional regulator YAP is a well-known mechanotransducer, that translocates to the nucleus upon increased cellular contractile forces, and extracellular stiffness. We therefore used YAP nuclear translocation as a positive control, as well as to optimise our screening protocol [10, 11]. We observed that at high cell density, where cells are known to show low actomyosin contractility, YAP was largely excluded from the nucleus. However, upon activation of RhoA, YAP was translocated to the nucleus (Fig. 1C and 1D). Using our biotinylation approach to purify nuclear proteins we reliably detected a more than 3-fold enrichment of YAP in the nucleus of tetracycline-, as compared to mock-treated cells (Fig. 1E, F, Supp. Fig. 2B, C). The presence of the nuclear marker nucleolin and the absence of the cytosolic marker tubulin confirmed the purity of our fractions. Together, this demonstrates that our overall screen approach can identify novel mechanotransducers that shuttle between the nucleus and cytoplasm in a mechanosensitive manner.

**Figure 1:**
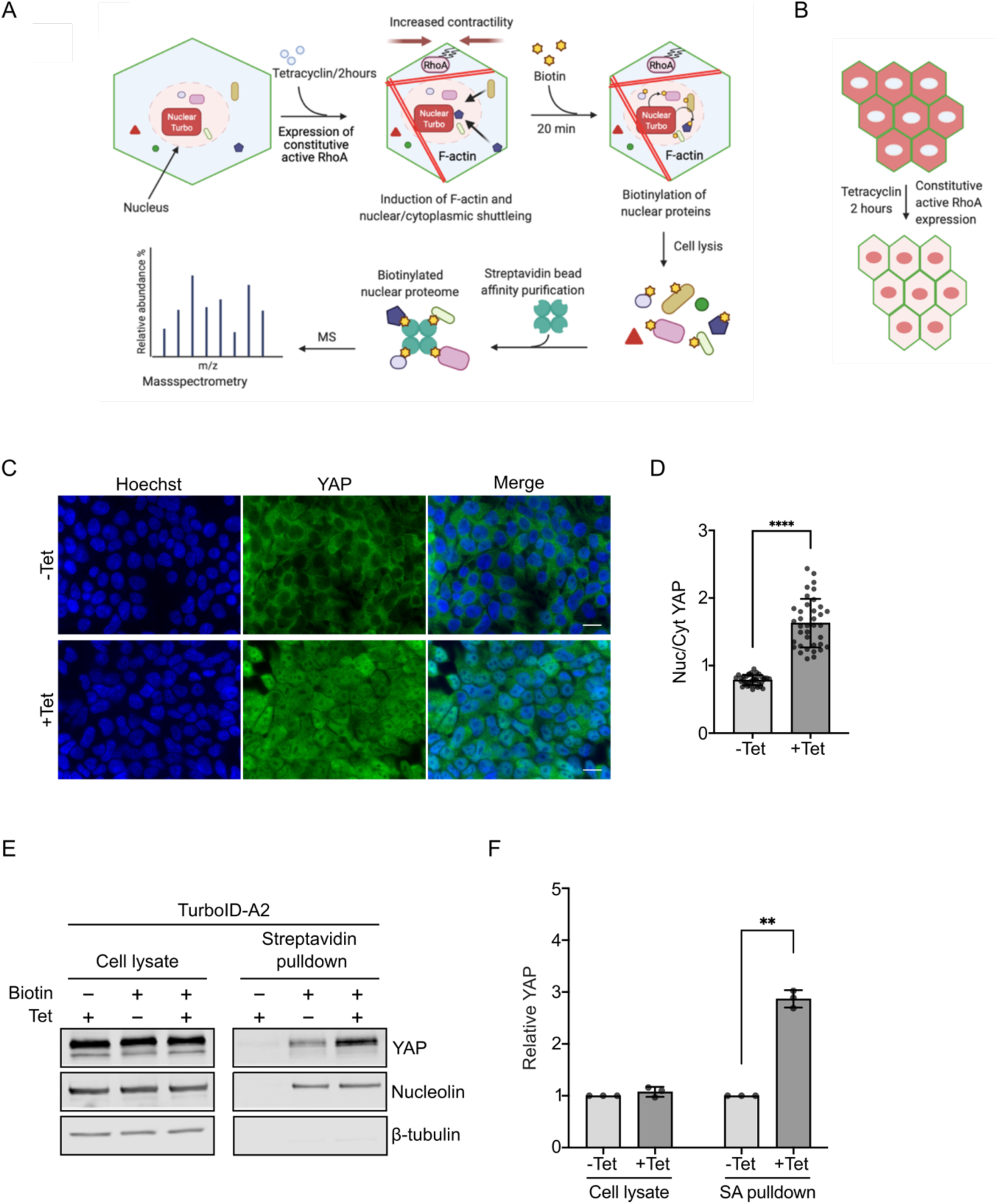
Screen approach to identify proteins potentially involved in mechanotransduction and its validation. **A:** Overview of the approach used for the screen. The diagram was created with BioRender.com. **B:** Scheme of nuclear translocation induced by tetracycline induced contractility at high cell density. The diagram was created with BioRender.com. **C-D:** Immunofluorescence and quantification of nuclear/cytoplasmic ratio of YAP. HEK cells stably expressing constitutive active RhoA under a tetracycline inducible promoter were treated for two hours with tetracycline. Cells were fixed and subcellular localisation was monitored by immunofluorescence. Bar represents 15 μm. Quantification shows the nuclear/cytosol ratio of YAP after tetracycline addition. Data was performed by unpaired t-test (n=3). Values are means ± s.d. \*\*\*\**p*<0.0001. **E-F:** Biochemical purification of nuclear proteins using the approach described in A. Biotinylated proteins were collected using streptavidin pulldown and probed by western-blotting using the indicated antibodies. Quantification shows normalised YAP expression Data was analysed by unpaired t-test (n=3). Values are means ± s.d. \*\**p*<0.01, ns: not significant.

### The splicing regulator PTBP1 is a potential mechanotransducer

To characterise protein abundance changes in the nuclear proteome after RhoA activation induced by tetracycline treatment, we measured the relative abundance of streptavidin-purified proteins using label-free quantitative mass spectrometry (Supplementary Table 1). Hierarchical cluster analysis of the quantitative mass spectrometry data showed that RhoA induction significantly changed the profiles of the proteins in the nucleus (Fig 2A). We identified 74 proteins that were enriched in the nucleus following RhoA induction, as described in the Methods and Material section (Fig. 2B). Importantly, our screen identified YAP with high confidence, which validated the overall approach. In addition to the proteins enriched in the nucleus upon tetracycline-treatment, we also found 543 proteins which showed lower amounts in the nucleus after addition of tetracycline using the same criteria as for the enriched proteins above. To validate the results from the mass spectrometry analysis, we quantified the nuclear enrichments of a few of the proteins identified, using nuclear fractionation prepared as described in the methods section followed by western-blot analysis. Thereby, we determined that the splicing regulator PTBP1, the molecular chaperon Chaperonin Containing TCP1 Subunit 2 (CCT2), and the transcription factor Paired Like Homeodomain 2 (Pitx2) showed nuclear enrichment after RhoA-activation (Fig. 2C). As expected, the positive control for the approach, YAP, also showed increased nuclear localisation upon activation of RhoA. Together, these data validate the mass spectrometry approach that we used (Fig. 2C). While the primary localisation of many of the proteins we identified to be enriched in the nucleus was known to be nuclear, we also found a significant number of proteins that were known to also localise to the cytosol or cytoplasm. (Fig. 2D). This is in line with the idea that proteins can shuttle between the nucleus and the cytoplasm upon a tetracycline-dependent increase in actomyosin contractility. The proteins upregulated in the nucleus were thereafter analysed with regard to a gene ontology, which showed an enrichment in proteins involved in RNA processing, RNA metabolic processes, and in transcriptional control (Fig. 2E, F). Specifically, proteins involved in the splicing of mRNA were enriched (Fig. 2E, Supplementary Table 2). This is in line with emerging evidence that specific alternative splicing events can be regulated by mechanical cues by mechanisms that are incompletely understood [25, 26]. Taken together, these findings validate our approach and demonstrate that RhoA-activation results in the nuclear enrichment of the splicing regulator PTBP1. Because PTBP1 has previously been reported to undergo shuttling between the nucleus and the cytoplasm, we decided to investigate further the role of PTBP1 in mechanotransduction [27].

**Figure 2:**
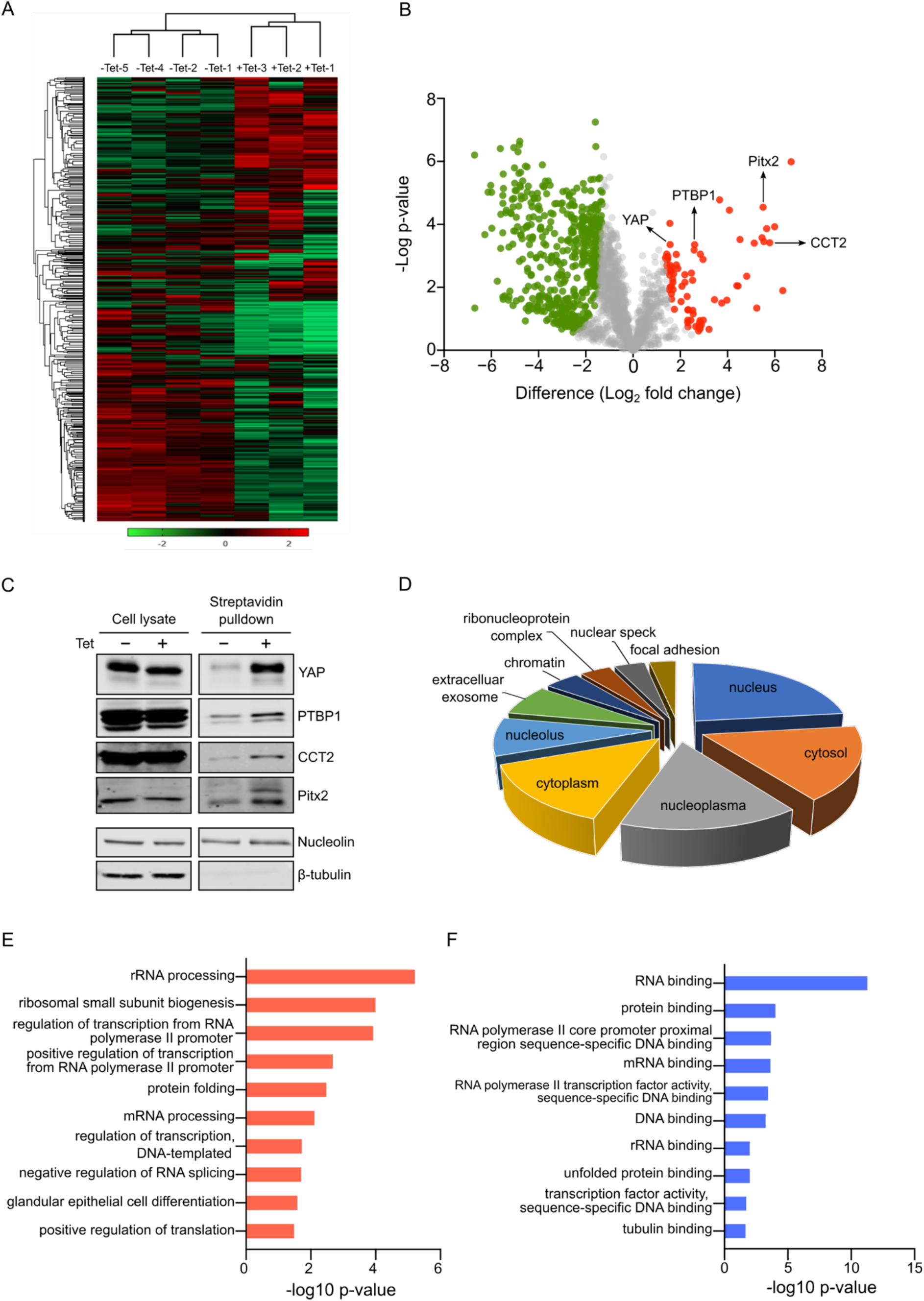
Nuclear proteome dynamics in response to constitutive active RhoA expression. **A**: Hierarchical cluster analysis of proteins showing increased or decreased abundance in the nucleus in HEK293-tet-RhoA-TurboID cells after 2 hours tetracycline treatment. Rows represent clustering of proteins and columns represent the clustering of samples. **B**: Volcano plot of proteins significantly enriched or depleted in the nucleus following the approach described in figure 1A and using the same data as used for the hierarchical cluster analysis. Red dots in the right area represent proteins significantly increased in the nucleus after tetracycline treatment while green dots indicate proteins significantly decreased in the nucleus by tetracycline treatment. **C**: Validation of upregulated candidates identified in the volcano-plot in B. Samples were prepared as described in 1A and probed by western-blotting using antibodies for the indicated proteins. **D**: Distribution of main subcellular localisations of proteins found significantly regulated in B. **E**: Gene ontology based on biological process (GOBP) analysis of proteome data showing the top 10 GO terms with significant enrichment after tetracycline treatment. **F**: Gene ontology based on molecular function (GOMF) analysis of proteome data revealed the top 10 most enriched GO terms after tetracycline treatment. All the proteome data were analysed by DAVID database (https://david.ncifcrf.gov/) and the criteria of differentially enriched nuclear proteins as permutation-based FDR < 0.05 with a Log2 nuclear enrichment > 1.5.

### Subcellular localisation of PTBP1 is regulated via cell density, ECM stiffness and actomyosin contractility

The screen used to identify novel mechanotransducers relies on the activation of RhoA, which, in addition to regulation of actomyosin contractility, also regulates other cellular signals and processes [28]. We therefore aimed to determine if the observed effects on the nuclear localisation of proteins was due to contractility-dependent effects of active RhoA. For this, we speculated that the nuclear localisation of proteins with a role in mechanotransduction would mimic the behaviour of the established mechanotransducer YAP. In cells that are cultured at a high cell density, or on an underlying soft extracellular matrix, YAP is excluded from the nucleus, while it is enriched in the nucleus at low cell density or in cells on a stiff extracellular matrix [10, 29]. Using normal, non-transformed MCF10A breast epithelial cells, we found that PTBP1 is excluded from the nucleus at high cell density while it is enriched in the nucleus at low cell density, a phenotype very similar to YAP (Fig. 3A). We validated the specificity of the antibodies against PTBP1 and YAP by siRNA that specifically targeted these proteins (Suppl. Fig. 3A, B, C). Furthermore, we also tested the subcellular localisation of PTBP1 on soft and stiff extracellular matrix (collagen-coated polyacrylamide gels of different stiffness) and found that PTBP1 is enriched in the cytosol on soft ECM but is enriched in the nucleus when cells are grown on stiff ECM (Fig.3B). We performed these experiments on custom-made as well as on commercial polyacrylamide gels, which were mechanically comparable (Suppl. Fig. 3D, E). Importantly, we also confirmed this finding using human primary mammary epithelial cells and in other cell lines (Suppl. Fig. 3F, G). Next, we investigated more directly whether the subcellular localisation of PTBP1 is regulated via actomyosin contractility, as has been previously described for YAP [10]. For this, NIH3T3 fibroblasts were cultivated on stiff conditions, on plastic, and incubated with different drugs that all inhibit actomyosin contractile forces: the actin polymerisation inhibitor latrunculin A, the myosin light chain kinase inhibitor ML-7, or the Rho kinase inhibitor Y27632. These inhibitors have previously been shown to reduce the nuclear localisation of YAP. As expected, all the inhibitors reduced the nuclear localisation, and increased the cytoplasmic localisation of YAP (Fig. 3C). Similarly, we observed a significant reduction of nuclear PTBP1, however, in contrast to YAP, we could not observe an enrichment of PTBP1 in the cytosol (Fig. 3C). We speculated that, given the brief time course of the experiment this was due to degradation of PTBP1 in the cytosol. To test this hypothesis, we blocked the proteasomal degradation in the presence of ML-7. Inhibition of protein degradation alone did not change subcellular localisation of PTBP1 or YAP, however, inhibition of proteasomal degradation in the presence of ML-7 led to a significant reduction of PTBP1 in the nucleus and an enrichment of PTBP1 in the cytosol. This suggests that after PTBP1 is translocated from the nucleus to the cytosol, it is rapidly degraded (Fig. 3D). Taken together, these data suggest that the subcellular localisation of PTBP1 is regulated by mechanical cues such as cell density and extracellular matrix stiffness, via a mechanism that requires actomyosin contractility. Thus, we conclude that the subcellular localisation of PTBP1 is under mechanical control, and therefore PTBP1 is a potential novel key protein in cellular mechanotransduction.

**Figure 3:**
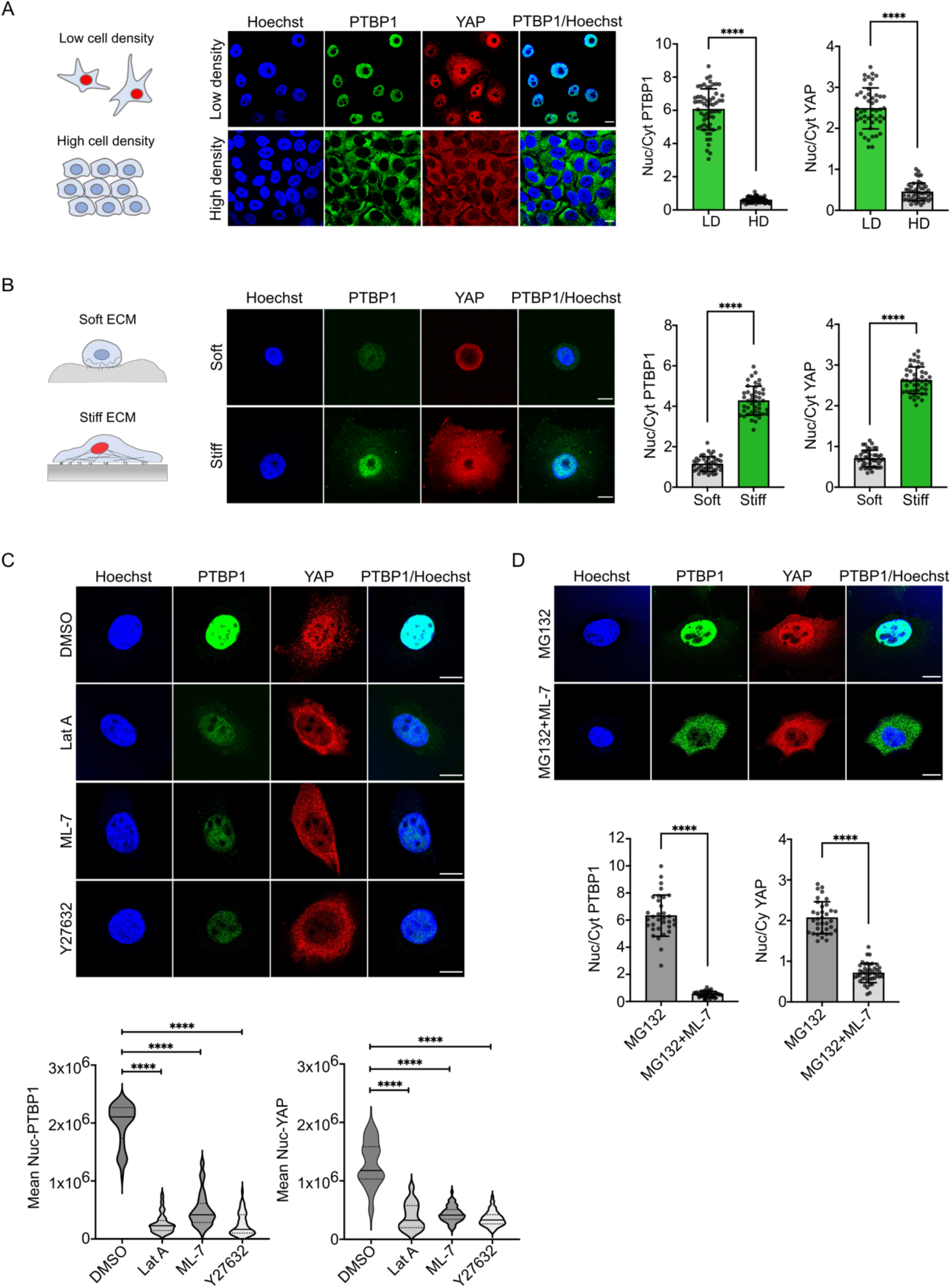
Subcellular localisation of PTBP1 is regulated by cell density, matrix stiffness and actomyosin contractility. **A**: Left and Middle: MCF10A cells were plated a high and low cell density and subcellular localisation of PTBP1 and YAP was determined by immunofluorescence using the indicated antibodies. Right: Quantification shows the nuclear to cytoplasmic ratio for PTBP1 and YAP at low cell density (LD) or high cell density (HD). Statistical analysis was performed by unpaired t-test. **B**: Left and Middle: MCF10A cells were plated on stiff (25kPa) or soft (0.2kPa) collagen coated polyacrylamide gels and subcellular localisation of PTBP1 and YAP was determined as described in A. Right: Quantification of nuclear to cytosolic ratio for PTBP1 and YAP on soft and stiff collagen-coated polyacrylamide gels. Data were analysed by unpaired t-test. **C**: Top: NIH3T3 fibroblasts were cultured on plastic and either treated with Latrunculin A (Lat.A), Myosin light chain kinase inhibitor-7 (ML-7) or the Rho kinase inhibitor Y27632 for 30 min. Cells were fixed and indicated proteins were identified by immunofluorescence. Bottom: Quantification of the mean nuclear intensities for PTBP1 and YAP after incubation with the inhibitors. Data were analysed by ordinary one-way ANOVA followed by Tukey’s multiple comparisons test. **D**: Top: NIH3T3 fibroblasts were incubated with the indicated inhibitors or combination of inhibitors and subcellular localisation of PTBP1 and YAP were determined using immunofluorescence. Bottom: Quantification of nuclear to cytosolic ratio for PTBP1 and YAP for cells treated with the proteasome inhibitor MG132 or treated with MG132 together with ML-7. Data were analysed by unpaired t-test. All experiments n=3, values are means ± s.d. \*\*\*\**p*<0.0001, ns: not significant. Bar in immunofluorescence images represents 10 μm.

### PTBP1 controls mechanically regulated splicing of the adapter protein Numb

To test the hypothesis that the mechanical control of the subcellular localisation of PTBP1 results in mechanically controlled splicing events, we investigated if the splicing of the well-established PTBP1 splicing target Numb depends on mechanical cues [30, 31]. Numb is an adapter protein that displays multiple functions, which include the regulation of cell fate choice [32], endocytosis [33, 34], and proliferation [35, 36]. Numb consists of an N-terminal Phosphotyrosine binding domain (PTB) and a c-terminal proline rich region (PRR). The Numb gene consists of 10 exons, and PTBP1 regulates the alternative splicing of exon 9, which results in the inclusion of exon 9 and the presence of an extra 46 amino acids in the proline rich region of the protein (Fig. 4A) [37, 38]. To test if the PTBP1-dependent alternative splicing of exon 9 of Numb is controlled by mechanical cues, we cultivated MCF10A breast epithelial cells on stiff or soft collagen-coated polyacrylamide gels and quantified the degree of inclusion of exon 9 in the mRNA, by polymerase chain reaction. We observed an increase of exon 9 inclusion in cells grown on stiff ECM as compared to cells grown on soft ECM (Fig. 4B). This is in line with our observation that a stiff underlying matrix increases the amount of PTBP1 in the nucleus. We further observed that the increase of this alternative splicing could be inhibited by siRNA that specifically targets PTBP1 (Fig. 4C). This indicates that PTBP1 is required for ECM stiffness dependent alternative splicing of the Numb mRNA. Other known PTBP1 targets such as the FAS receptor or cortactin (CTTN) showed similar matrix stiffness dependent alternative splicing dependent on PTBP1 (Suppl. Fig. 4A-D). Next, we asked the question whether the splicing change observed at the mRNA level for Numb can also be observed at the protein level. MCF10A cells were cultured on soft and stiff ECM and Numb isoform expression was quantified using western-blotting. We observed two variants of Numb, which appeared as a duplet band, where the upper band matches the size of the protein isoform(s) which includes the additional 46 amino acids coded for by exon 9. Stiff culture conditions increased the longer splicing isoform of Numb, with an increase of the ratio of the long variant of Numb with exon 9 versus the shorter variant without exon 9 from 1.2 to 3 (Fig. 4D). Thus, we conclude that ECM stiffness can regulate the ratio of Numb protein isoforms increasing the longer splicing variant over the shorter one.

**Figure 4:**
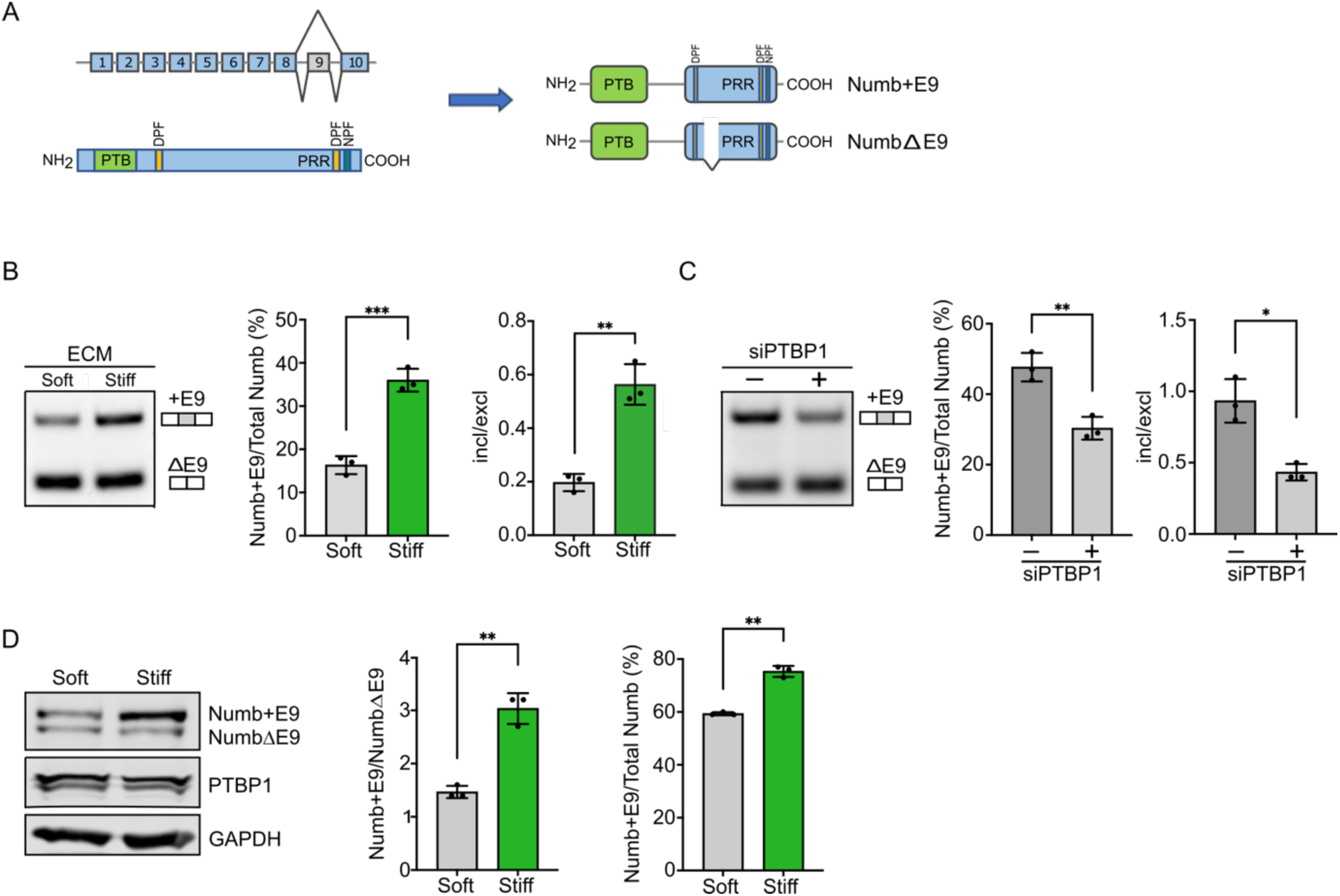
PTBP1 regulates matrix stiffness dependent splicing of Numb. **A**: top left: Exon-intron structure of Numb. Top bottom: Modular structure of the Numb protein, PTB= phosphotyrosine binding domain, PRR= proline rich region. Right: Modular structure of Numb showing the localisation of the 49 amino acid insert encoded by exon 9. **B**: MCF10A cells were cultured for 5 days on soft PAA gels and then cultured for 3 days on soft or stiff PAA gels. Exon 9 inclusion (+E9) and the ratio of inclusion versus exclusion (ΔE9) was measured via reverse transcription followed by PCR. The relative exon 9 (%) was defined as the ratio of exon 9-included isoform over the total level of exon 9-included and exon 9-excluded isoforms, while the incl/excl was calculated as the ratio of exon 9-included isoform against exon 9-excluded isoform. **C**: MCF10A cells were cultured on plastic and transfected with the siRNA targeting PTBP1 or control siRNA. Exon 9 inclusion and inclusion versus exclusion ratio were determined as described in B. **D**: MCF10A cells were incubated on soft polyacrylamide gels for 5 days and then split on soft and stiff polyacrylamide gels. Total protein was isolated after 5 days and analysed by western-blotting using the indicated antibodies. Blot of GAPDH was used as loading control. Exon 9 inclusion and inclusion versus exclusion ratio was determined by densitometric quantification of corresponding western-blots. All experiments n=3. Data were analysed by unpaired t-test. Values are means ± s.d. \**p*< 0.05, \*\**p*< 0.01, \*\*\**p*< 0.001.

### Alternative splicing of Numb is required for stiffness-induced cell proliferation

The stiffness of the extracellular matrix is an important regulator of cell proliferation, however, the underpinning mechanisms are not fully understood [39, 40]. In general, increased stiffness of the ECM increases cell proliferation. The function of Numb isoforms has been investigated widely, and splicing variants which include exon 9 promote proliferation, whereas variants without exon 9 inhibiting or do not change cell proliferation [35, 41]. Therefore, we speculated that PTBP1 in promoting the exon 9 including Numb splicing isoform on a stiff matrix might contribute to stiffness-induced proliferation. To test this, we analysed the cell proliferation of MCF10A breast epithelial cells on soft or stiff collagen-coated polyacrylamide gels. As expected, cell proliferation was significantly higher on a stiff matrix as compared to a soft matrix. While the knockdown of PTBP1 did not change the proliferation on soft ECM, it significantly reduced the proliferation on stiff ECM, to almost the level observed on soft matrices (Fig. 5A). This suggests that PTBP1 is required for extracellular stiffness-induced cell proliferation. To further investigate the molecular mechanism of this control, we next examined if overexpression of Numb isoforms can rescue the reduced proliferation phenotype on stiff ECMs observed upon loss of PTBP1. For this, we transfected MCF10A cells with siRNA targeting PTBP1 followed by a transfection of the Numb isoform containing exon 9 (+E9) or the Numb isoform that lacks exon 9 (ΔE9). PTBP1 knockdown was confirmed by immunofluorescence and successful overexpression of Numb isoforms was confirmed by western-blotting, as shown in the figure 5 (Fig. 5B, C). We observed that expression of the Numb isoform with but not the isoform without exon 9 rescued the proliferation rates of PTBP1 depleted MCF10A cells on stiff ECM to almost normal proliferation rates (Fig. 5B). These findings are in line with the hypothesis that extracellular-stiffness-induced cell proliferation depends on PTBP1 promoting Numb exon 9 inclusion on stiff ECM. To further test this hypothesis, we reduced the levels of endogenous Numb isoforms, using isoform-specific siRNAs. For this, MCF10A cells were cultured on stiff ECM and transfected with siRNA targeting either the +E9 isoform or the ΔE9 isoform. In line with our working hypothesis, we observed that only downregulation of the +E9 isoform reduced cell proliferation but not downregulation of the ΔE9 isoform (Fig. 5D, E). Finally, we tested whether overexpression of Numb isoforms would affect proliferation on soft ECM. We observed that while the overexpression of the ΔE9 isoform did not have any effect on the proliferation on soft ECM, the overexpression of +E9 isoform allowed proliferation on soft ECM, even to levels that were comparable to the proliferation levels observed on stiff ECM (Fig. 5F, G). We found similar results using MDCK cells that overexpress the +E9 isoform under a tetracycline inducible promoter but not with MDCK cells overexpressing the ΔE9 isoform (Suppl. Fig. S5 A-F). Therefore, overexpression of the Numb splicing variant that includes exon 9 is sufficient to override the inhibition of proliferation in cells on soft ECM. In summary, our data suggest that PTBP1-regulated alternative splicing of Numb is crucial for extracellular stiffness-induced cell proliferation.

**Figure 5:**
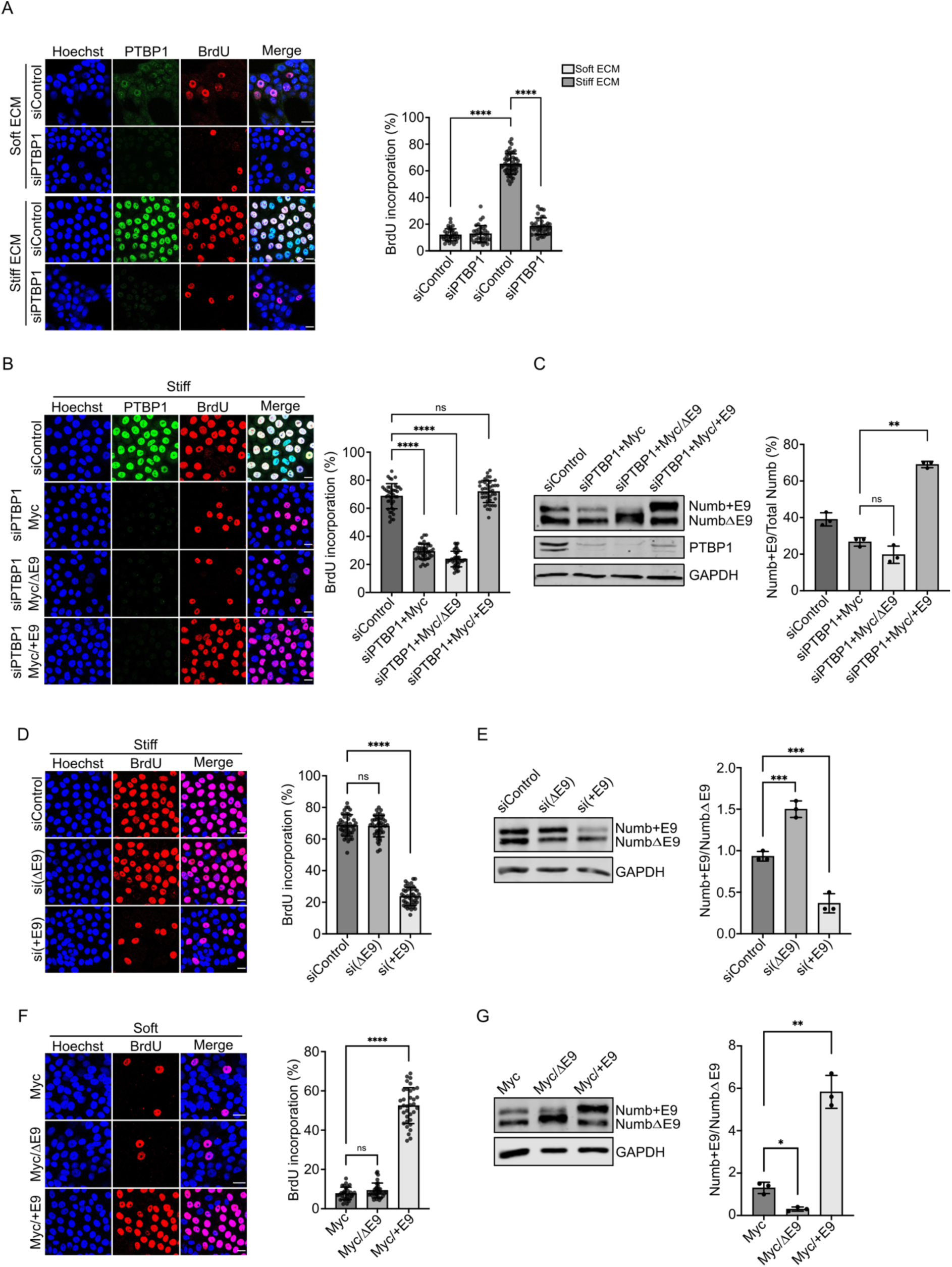
PTBP1 and Numb splicing is important for matrix stiffness induced proliferation. **A**: Right: MCF10A cells were cultured on stiff or soft collagen coated polyacrylamide gels and transfected with the indicated siRNAs. Proliferation rate and knockdown was determined by immunofluorescence. Left: Bar diagram shows the corresponding quantification of BrdU incorporation. **B**: Left: MCF10A cells were cultured on a soft or stiff matrix and transfected with the indicated siRNAs and Numb isoform specific expression constructs Myc+E9 and Myc/ΔE9. Right: The proliferation rate was quantified using BrdU incorporation. **C:** Western-Blot showing successful knockdown of PTBP1 and overexpression of Numb isoforms. Numb isoforms were detected using an anti-Numb antibody and signals represent the sum of endogenous Numb and the respective overexpressed isoform. Bar diagram shows quantification of Numb isoform overexpression as percentage of Numb+E9 of total Numb expression. **D**: MCF10A cells were grown on stiff collagen coated polyacrylamide gels and transfected with the indicated isoform specific siRNAs si(+E9) and si(ΔE9) and proliferation rate was measured using a BrdU assay. Bar diagram represents quantification of BrdU incorporation. **E**: Western blot showing successful knockdown of endogenous Numb isoforms. Bar diagram shows quantification of knockdown as Numb+E9 to NumbΔE9 ratio. **F:** MCF10A cells were cultured on soft collagen coated polyacrylamide gels and transfected with the indicated isoform specific Numb expression constructs. Proliferation rate was measured and quantified by BrdU assays. **G:** Western blot validating overexpression of Numb isoforms. Bar diagram shows quantification of Numb isoform overexpression as a change in the Numb+E9 to NumbΔE9 ratio. All experiments n=3. Statistical analysis was performed by Data were analysed by ordinary one-way ANOVA followed by Tukey’s multiple comparisons test. Values are means ± s.d. \*\*\*\**p*<0.0001. Bar in immunofluorescence images represents 20 μm.

### Numb +E9 isoform is important for mechanomemory

We have demonstrated that extracellular stiffness-induced regulation of PTBP1 results in stiffness-dependent changes of the expression of Numb isoforms. This opens the possibility that these changes contribute to the formation of a mechanomemory. Mechanomemory or mechanical conditioning of cells describes the observation that cells retain, for some time behaviour acquired in their previous mechanical environment, when placed in a different mechanical environment [42, 43]. We tested whether MCF10A cells can form a mechanomemory with respect to stiffness-induced cell proliferation, by first culturing the cells on stiff ECM or soft ECM, followed by replating them on an underlying matrix of different stiffness, followed by quantification of their proliferation rate (Fig. 6A). We observed that cells pre-cultured on stiff ECM showed a much higher (5.6-fold) proliferation rate on soft ECM, than cells pre-cultured on soft ECM. Similarly, cells previously grown on soft ECM showed a slower (0.7-fold) proliferation rate on stiff ECM than cells previously grown on stiff ECM (Fig. 6B). To determine if the increased proliferation rate on soft ECM of cells previously grown on stiff ECM depends on increased expression levels of the +E9 Numb isoform, cells were grown on stiff ECM and then transfected with siRNA targeting the Numb isoforms with or without exon 9. Cells were then re-plated on soft ECM and cell proliferation quantified (Fig. 6C). Knockdown of the Numb ΔE9 isoform did not change cell proliferation, however, knockdown of the +E9 Numb isoform almost completely abolished the memory effect (Fig.6D). In summary, our data are in line with the idea that upregulation of the Numb isoform that includes exon 9 is part of the mechanism that underpins the formation of a mechanomemory for cell proliferation.

**Figure 6:**
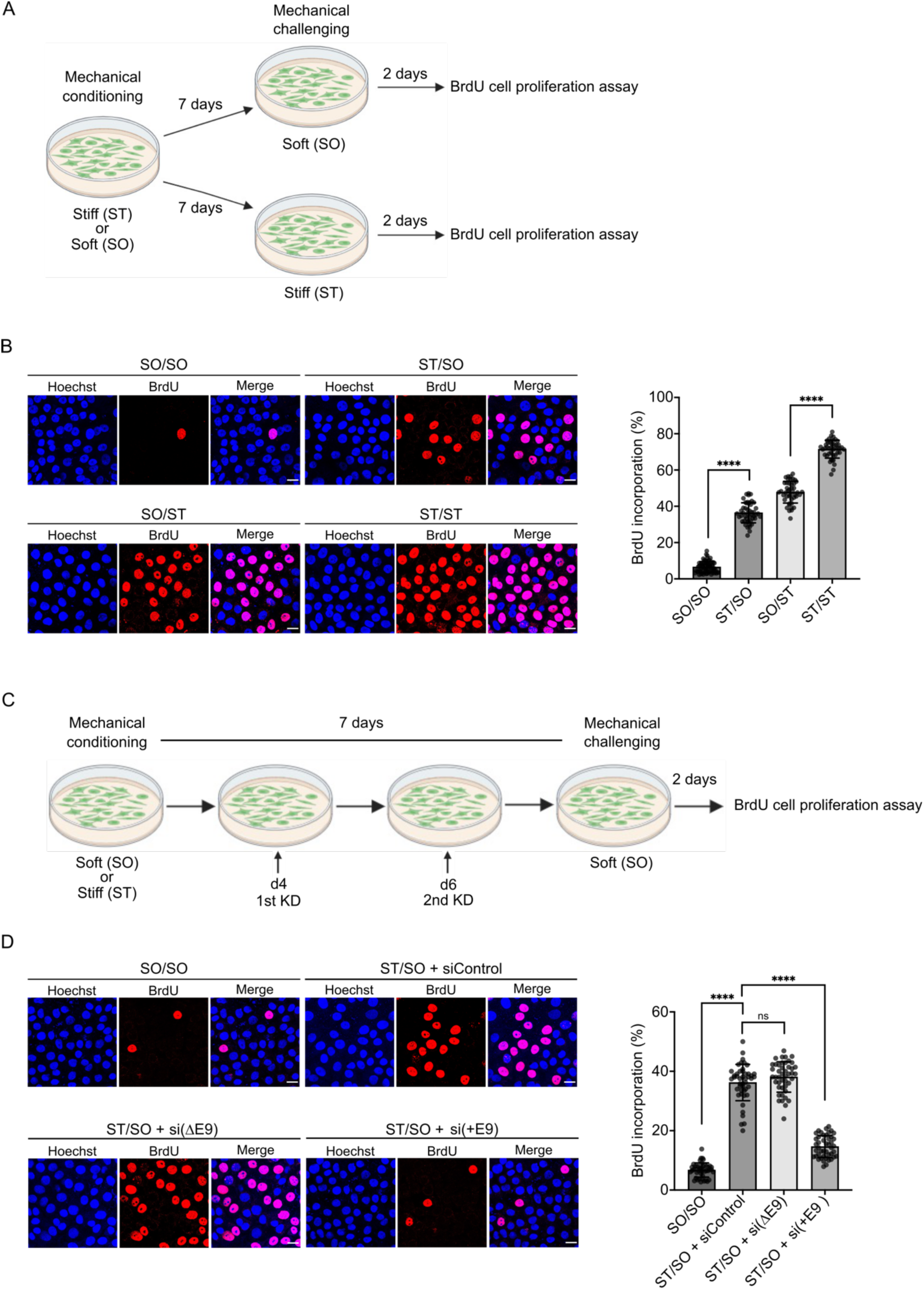
Mechanosensitive Numb splicing is important for mechanomemory formation. **A**: Protocol to test for capacity to form a mechanomemory. MCF10A cells were cultured on soft or stiff collagen coated polyacrylamide gels for 7 days (mechanical conditioning phase) and then replated on stiff or soft polyacrylamide gels (mechanical memory test phase) for 2 days and followed by a BrdU assay to measure proliferation. The diagram was created with BioRender.com. **B**: BrdU assay data following the protocol described in A. Bar diagram shows quantification of the BrdU assay. **C**: Protocol to test involvement of Numb isoform expression in mechanomemory formation. MCF10A cells were cultured on stiff collagen coated polyacrylamide gels for 7 days. Cells were transfected with control or isoform specific NUMB siRNAs si(+E9) and si(ΔE9) at day 4 and day 6. Cells were then replated on soft polyacrylamide gels and 2 days later a BrdU assay performed. The diagram was created with BioRender.com. **D**: BrdU assay data following the protocol described in C. Bar diagram shows quantification of BrdU assays. All experiments n=3. Statistical analysis was performed by unpaired t-test. Values are means ± s.d. \*\*\*\**p*<0.0001,

## DISCUSSION

Here, we present a novel proteomic-based screen that allows the identification of proteins involved in mechanotransduction. Our screen identified the known, and well-studied mechanosensitive transcriptional regulator YAP, which demonstrates the suitability of our screen to identify proteins that mediate mechanotransduction. Using this screen, we found that the subcellular localisation of the splicing regulator PTBP1 is mechanosensitive, a finding that suggested that PTBP1 can play a role in mechanotransduction. In line with this concept, emerging evidence suggests that extracellular mechanical and physical cues can control alternative splicing. Changes in alternative splicing have been described after mechanical loading of bone or after stretching of cells [44, 45]. Tissue stiffness has also been shown to regulate serine/arginine rich (SR) splicing factors, which leads to stiffness-dependent splicing of the extra domain B-fibronectin isoform which likely plays a role in tumour progression [25]. Similarly, matrix stiffening reduces the expression levels of the epithelial splicing regulatory protein Epithelial splicing regulator (ESPR1), which regulates alternative splicing of the regulator of actin dynamics MENA, and thereby tumour cell intravasation [46]. Thus, the mechanical regulation of alternative splicing appears to be an important principle to translate mechanical cues into cellular biochemical changes, to govern mechanosensitive cell behaviour. However, the investigation of alternative splicing as an integral part of mechanotransduction is in its infancy. Here, we add the splicing regulator PTBP1 among the mechanosensitive splicing factors, and demonstrate that actomyosin contractility directly controls the subcellular localisation of this splicing factor, a principle previously demonstrated for transcriptional regulators such as YAP and MRTF-A [10, 12]. It has recently been demonstrated that direct application of physical force to the nucleus modulates nuclear pore permeability and affects facilitated diffusion. Furthermore, forces can be transmitted to the nucleus via the actomyosin cytoskeleton and the LINC complex leading to nuclear pore deformation and changes in permeability [4, 13]. It is known that cells display increased actomyosin contractile force when cultured on a stiff matrix [47]. We show that lowering cellular actomyosin contractility via small molecule intervention or by placing cells on a soft matrix, reduces the nuclear localisation of PTBP1. Hence, it is likely that a mechanically controlled nuclear permeability or mechanically controlled facilitated diffusion might contribute to the regulation of the subcellular localisation of PTBP1.

To investigate a role of PTBP1 in mechanotransduction we have investigated the potential mechanosensitive regulation of PTBP1 splicing targets. We found that PTBP1 regulates the matrix stiffness-dependent splicing of the adapter protein Numb among other splicing targets, and that this is accompanied by a change in protein expression of the corresponding Numb isoforms. Numb plays important roles in cell fate decisions, regulation of endocytic trafficking and lysosomal degradation of membrane receptors including Notch1, E-cadherin and the anaplastic lymphoma kinase [48, 49]. Numb not only undergoes alternative splicing of exon 9 but also exon 3, giving rise to four different isoforms in vertebrates. Exon 9 encodes for an additional 49 amino acids in the proline rich region of Numb and gives rise to the two largest isoforms P71/P72. Exon 3 codes for an 11 amino acid insert into the PTB domain of Numb [37]. The splicing of Numb is regulated during development with the isoforms that include exon 9 expressed in stem and progenitor cells and the isoform skipping exon 9 preferentially expressed in differentiated cells [50, 51]. These isoforms have been described to serve different and partially opposite cellular functions [35, 50]. Furthermore, Numb exon 9 inclusion is increased in multiple cancers including all breast cancer subtypes. This is of particular interest given that isoforms including exon 9 promote proliferation which can contribute to cancer progression [30, 41, 52, 53]. Similarly, mutations in the splicing factor RBM10 lead to increased exon 9 inclusion in Numb which promotes lung cancer cell growth [35]. Although the role of Numb isoforms in cancer is complex and appears to be cell type dependent [54, 55], this indicates that our finding can be important for cancer progression.

Here, we demonstrate that mechanically controlled Numb splicing is crucial for matrix stiffness-induced proliferation. Matrix stiffness can control cell proliferation via multiple mechanisms, which include stiffness-dependent activation of ERK kinases, AKT and the signal transducer and activator of transcription (STAT) -3. In breast epithelial cells, matrix stiffness increases cell proliferation via the oncogene ZNF217, and increased levels of ZNF217 are found in cells from dense breast tissue, which is linked to increased lifetime risk for breast cancer [56]. Furthermore, cells tend to spread more on a stiff underlying matrix as compared to on a soft matrix, which is at least in part driven by increased force over cell-matrix adhesions, and by an increase in stress fibres [47, 57]. Cell spreading can positively regulate cell proliferation [58, 59]. Finally, as described above, ECM stiffness drives the nuclear translocation of the transcriptional regulator YAP, which regulates transcription of cell cycle regulators and genes directly involved in the regulation of DNA synthesis, repair and mitosis, which promotes cell proliferation [60–63]. The mechanism of how the Numb +E9 isoform promotes stiffness-induced cell proliferation is currently unknown. However, because Numb regulates receptor trafficking [37], it is possible that it results in increased recycling of receptor tyrosine kinases that are important for proliferation, which would increase the signalling input for cell proliferation in cells subjected to a stiff environment. This is supported by the fact that the Numb +E9 isoform but not the ΔE9 isoform regulates the recycling of the receptor tyrosine kinase ALK [64]. Furthermore, a recent study has shown that the Numb +E9 isoform is important for the cell surface expression of several receptors, including integrins, which can significantly increase the spreading of cells on a stiff matrix, which could contribute to the observed increase in cell proliferation [52, 58, 59].

We also tested the involvement of Numb alternative splicing in the formation of a mechanomemory. Mechanomemory can act on different time scales from seconds to several days and can be reversible, partially reversible, or irreversible [65]. The molecular mechanisms underpinning mechanomemory formation are currently only emerging. This includes for example the mechanical control of chromatin condensation, which appears to play a role in irreversible differentiation of fibroblasts into myofibroblasts [66]. Also, mesenchymal stem cells cultured on a stiff matrix for prolonged times show a gradual increase in the expression of the microRNA-21, and this expression remains high even after switching cells to a soft environment, which renders the cells insensitive to subsequent exposure to soft matrix and induces a pro-fibrotic cell programme [67]. Here we demonstrate that in addition to these mechanisms, also changes in alternative splicing of mRNA can contribute to the formation of a mechanical memory for cell proliferation. We expect that the duration of this memory depends on the time scale of the reversibility of the nuclear localisation of the splicing factor, as well as the half-lives of the protein isoforms and of their corresponding isoform-specific RNAs. In summary, our work has identified a novel mechanism by which matrix stiffness can promote cell proliferation, and that this mechanism can also contribute to a mechanomemory in cells. Our results have implications for tissue development and for diseases like cancer and fibrosis, which show higher levels of tissue stiffness than healthy tissue [68]. Targeting alternative splicing using antisense oligonucleotides is already a clinical reality and further knowledge of mechanically regulated splicing events in disease context will allow to develop novel cancer or fibrosis mitigating strategies [69].

## LIMITATIONS

It is unlikely that our screen was exhaustive given the fact that we did not find known mechanosensitive transcription factors like MRTF-A or Twist. This might be partially due to the cell type (HEK cells) the initial screen was performed in. Our findings show that PTBP1, via the regulation of alternative splicing of Numb, regulates stiffness-induced cell proliferation. However, further studies are required to identify the exact underpinning mechanism. Furthermore, we have confirmed matrix stiffness-dependent subcellular localisation of PTBP1 in primary human mammary epithelial cells but much of our findings relies on the non-transformed breast epithelial cell line MCF10A, it will be important to investigate the contribution of the proposed mechanism using primary epithelial cells or within tissue context.

## ACKNOWLEDGMENTS

We are very grateful to Sanjay Kumar and Stacey Lee, University of California, Berkeley, United States, and Manuel Théry and Benoit Vianay, Institut de Recherche Saint Louis, France, for help and advice on traction force microscopy analysis. We acknowledge funding from the Libyan government for A.S.

## DATA AVAILABILITY STATEMENT

The mass spectrometry proteomics data have been deposited to the ProteomeXchange Consortium via the PRIDE partner repository with the dataset identifier PXD046157. All other data supporting the findings of this study are available from the corresponding author on reasonable request.

## AUTHOR CONTRIBUTIONS

P.L.T and K.S.E conceived the study. P.L.T., K.S.E., A.K.B.G, M.O.C. designed the experiments. P.L.T., W.S., A.S., S.M., M.O.C. performed the experiments. S.M. and A.S. performed the traction force microscopy, M.O.C. performed the mass spectrometry experiments and analysed the data, all other experiments were performed by P.L.T. and W.S.. A.K.B.G and S.M. analysed the traction force microscopy data. K.S.E and P.L.T. wrote the manuscript. All authors commented on the manuscript with special input from A.K.B.G.

## Method details

### Cell lines

Flp-In ^TM^T-REX^TM^ HEK293 were cultured with DMEM-GlutaMAX^TM^ media (Gibco) supplemented with 10% FBS (Gibco), 1 % penicillin/streptomycin (Gibco), 100 μg/ml zeocin (Gibco) and 15 μg/ml blasticidin S HCl (Gibco). Flp-In TREX-MDCK II cells (gift from Dr Jack Kaplan, University of Illinois at Chicago) were cultured with DMEM-GlutaMax^TM^ supplemented with 10% FBS, 1 % penicillin/streptomycin, 150 μg/ml zeocin and 6 μg/ml blasticidin S/HCl. MCF10A (ATCC) were cultured with DMEM/F12 media (Gibco) supplemented with 5% Horse Serum, 20 ng/ml EGF (Sigma-Aldrich), 500 μg/ml hydrocortisone (Sigma-Aldrich), 10 μg/ml insulin (Thermo Fisher Scientific)) and 1 % penicillin/streptomycin (Gibco). NIH-3T3 cells (ATCC) were cultured with DMEM-GlutaMax^TM^ supplemented with 10% FBS and 1 % penicillin/streptomycin. Human mammary epithelial cells (Sigma-Aldrich) were cultured with Human Mammary Epithelial Cell Growth Medium (Sigma-Aldrich). HeLa (ATCC) were cultured with DMEM-GlutaMax^TM^ supplemented with 10% FBS and 1 % penicillin/streptomycin. Cells were grown at 37°C in a humidified atmosphere containing 5% CO2, and cells of early passages were used for the experiments.

### Vector constructs

pcDNA3-EGFP-RhoA-Q63L was a gift from Gary Bokoch (Addgene plasmid # 12968) and 3xHA-TurboID-NLS_pCDNA3 was a gift from Alice Ting (Addgene plasmid # 107171). Numb+E9_pcDNA3.1(+)-C-Myc and Numb⊗E9_pcDNA3.1(+)-C-Myc isoforms were purchased from GenScript. pcDNA^TM^5/FRT/TO and pOG44 vectors were from Thermo Fisher Scientific). pcDNA3.1 (+)-myc-PuroR was generated in Dr Kai Erdmann Lab (University of Sheffield, UK).

To generate Tet-induceable constitutively active RhoA (CA-RhoA) expression vector, the full-length cDNA encoding EGFP-RhoA-Q63L was amplified from pcDNA3-EGFP-RhoA-Q63L by PCR, with primers Q63L-RhoA-F (5’-CTTAAGCTTGCCATGGTGAGCAAGGGC-3’) and Q63L-RhoA-R (5’-CTTGCGGCCGCTCACAAGACAAGGCAACCAGATTTTT-3’), using the Phusion^TM^ High-Fidelity Polymerase (Thermo Fisher Scientific) according to manufacturer’s instructions. The EGFP-RhoA-Q63L amplicon was purified and then ligated into the pcDNA^TM^5/FRT/TO vector using HindIII and NotI sites to form the plasmid pcDNA5/FRT/TO-RhoA.

To generate TurboID expression vector pcDNA3.1-Puro-3xHA-TurboID-NLS for proximity labelling of nuclear proteomes, the cDNA region encoding HA-tagged TurboID-NLS was prepared from 3xHA-TurboID-NLS_pCDNA3 using endonuclease digestion and then integrated into pcDNA3.1 (+)-myc-PuroR vector using HindIII and XbaI restriction sites.

To generate plasmids expressing Numb+E9 or Numb⊗E9 specific isoforms under the control of a tetracycline inducible promoter, cDNA region encoding Numb+E9 or Numb⊗E9 were amplified from either Numb+E9_pcDNA3.1(+)-C-Myc or Numb⊗E9_pcDNA3.1(+)-C-Myc by PCR with the primers FRT/TO-Numb-F (5’-CCCAAGCTTGCCACCATGAACAAATTACGGCAAAGT-3’) and FRT/TO-Numb-R (5’-CGCCTCGAGTCACAGATCCTCTTCAGAGATGAGTTTCT-3’), using Phusion^TM^ High-Fidelity Polymerase (Thermo Fisher Scientific) according to manufacturer’s instructions. The purified PCR fragments were ligated into pcDNA^TM^5/FRT/TO vector using HindIII and XhoI sites to create the plasmids pcDNA5/FRT/TO-Numb+E9 and pcDNA5/FRT/TO-Numb⊗E9.

### siRNA transfections

Gene knockdown experiments were carried out by siRNA transfections with Lipofectamine^TM^2000 (Thermo Fisher Scientific) according to the manufacturer’s instructions. siRNA and stealth RNAi negative control used in this study are listed in the Key Resources Table. In brief, 4x10^5 MCF10A cells were seeded in 35mm dish for 24 hours followed by transfection with a preincubated mixture of 5 pmole siRNA, 125 μl of Opti-MEM, and 5 μl of Lipofectamine 2000 for 6 hours. The medium was replaced, and cells were allowed to recover overnight before second siRNA transfection was performed. Knockdown efficiency was verified by western blotting or immunofluorescence staining for the targeted proteins.

### Generation of stable cell lines

To create HEK293-tet-RhoA-TurboID stable cell lines harbouring tetracycline-inducible constitutively active RhoA and TurboID, pcDNA5/FRT/TO-RhoA-Q63L and pcDNA3.1-Puro-3xHA-TurboID-NLS were successively introduced into Flp-In^TM^ T-REX^TM^ HEK293 cells. In brief, Flp-In^TM^ T-REX^TM^ HEK293 cells were seeded in 6-well plates and allowed to grow to about 70% confluency prior to co-transfection of pcDNA5/FRT/TO-RhoA-Q63L and pOG44 using Lipofectamine^TM^2000 (Thermo Fisher Scientific) according to the manufacturer’s instructions. 24 hours after transfection, the medium was changed to complete medium (DMEM-GlutaMAX^TM^ containing 15 μg/ml of blasticidin S HCl) for another 24 hours. Transfected cells were then cultured with selection media (DMEM-GlutaMAX^TM^ supplemented with 15 μg/ml of blasticidin S/HCl, and 200 μg/ml of hygromycin B) until the cell colonies (named HEK293-tet-RhoA) were formed and collected for subsequent transfection with linearised pcDNA3.1-Puro-3xHA-TurboID-NLS. After 24 hours, transfected cells were split in a 1:10 ratio and cultured with selection media (DMEM-GlutaMAX^TM^ supplemented with 15 μg/ml of blasticidin S/HCl, 200 μg/ml of hygromycin B, and 1.5 μg/ml of puromycin). Single colonies were picked and expanded. Only clones exhibiting Tet-inducible CA-RhoA and biotinylation activity were selected for further experiments.

To generate MDCK/Numb+E9 and MDCK/Numb⊗E9 stable cell lines, pcDNA5/FRT/TO-Numb+E9 or pcDNA5/FRT/TO-Numb⊗E9 was co-transfected with pOG44 into Flp-In T-REX MDCK using Lipofectamine^TM^2000 (Thermo Fisher Scientific) according to the manufacturer’s instructions. Stable clones were selected using selection media (DMEM-GlutaMAX^TM^ supplemented with 6 μg/ml of blasticidin S/HCl, and 150 μg/ml of hygromycin B) until single colonies were formed and collected.

### Western blotting

Standard western blotting techniques were used, and samples were kept on ice during the experimental procedure. Cells were lysed in ice-cold RIPA buffer (50 mM Tris-HCl pH 7.5, 150 mM NaCl, 0.1% (w/v) SDS, 0.5% (w/v) sodium deoxycholate, 1 mM EDTA, 1% (v/v) NP-40) supplemented with protease cocktail inhibitor and PhoSTOP^TM^ phosphatase inhibitor (Sigma-Aldrich). The cell lysate was centrifuged at 16600 x g for 20 minutes at 4°C and the supernatant was collected for western blotting. Protein concentration was determined using Bio-Rad DC protein assay kit according to manufacturer’s instructions. Equal amounts of total protein were loaded onto an SDS-PAGE before transfer onto a nitrocellulose membrane (Millipore). The membranes were blocked in 5% (w/v) milk in PBST (0.05% Tween 20, 1xPBS) for 1 hour, followed by incubation overnight at 4°C with primary antibodies diluted in 5% (w/v) milk in PBST or in 5% (w/v) BSA in TBST (13.7 mM NaCl, 2 mM Tris-HCl, 0.1% Tween 20 pH 7.6) and then 1 hour at room temperature with secondary antibodies. Primary antibodies were used at the following dilutions: YAP 1:200 (sc-101199; Santa Cruz), GFP 1:2000 (A-11122; Invitrogen), PTBP1 1:500 (32-4800; Thermo Fisher Scientific), Pitx2 1:1000 (PA-1020; Capra Science), CCT2 1:200 (sc-374152; Santa Cruz), Nucleolin 1:300 (sc-8031; Santa Cruz), Numb 1:1000 (2756; Cell Signalling Technology), β-tubulin 1:5000 (T4026; Sigma-Aldrich), γ-adaptin 1:1000 (gift from Dr Andrew Peden, University of Sheffield), GAPDH 1:20000 (60004-1-IG; Proteintech), beta-catenin 1:2000 (610153; BD Biosciences). Donkey anti-rabbit IgG (H+L) Alexa Fluor^®^680 (A21109; Thermo Fisher Scientific) and DyLight^TM^800 4x PEG conjugate anti-mouse IgG (H+L) (SA5-35521; Thermo Fisher Scientific) were used as secondary antibodies for infrared detection. Results were visualized and quantified with Odyssey Sa^®^ Infrared imaging system (LI-COR).

### Nuclear protein biotinylation and LC-MS/MS analysis

HEK293-tet-RhoA-TurboID were grown at high cell density before treatment with 500 μM biotin for 20 minutes at 37°C to allow the biotinylation of the nuclear proteomes. Cells were placed on ice to stop labelling followed by washing with ice-cold PBS to remove excess biotin. Cells were then lysed in ice-cold RIPA buffer supplemented with protease cocktail inhibitor and PhoSTOP^TM^ phosphatase inhibitor (Sigma-Aldrich). The cell lysate was centrifuged at 16600 x g for 20 minutes at 4°C and the supernatant was collected. Cell lysates containing 3000 μg protein were then incubated overnight with 100 μl of Pierce^®^ Streptavidin Agarose Resins (Thermo Fisher Scientific) at 4°C. The mixture of cell lysate/streptavidin agarose was transferred to a Wizard^R^ Minicolumn (Promega) and the resins were washed with buffers in the following order: 10 ml of 2% SDS, 10 ml of RIPA buffer, 10 ml of 0.5 M NaCl, 10 ml of 2M Urea/50 mM Tris-HCl pH8, and 10 ml of 50 mM ammonium bicarbonate. The resins were then transferred to another clean eppendorf and washed twice with 1 ml of 50 mM ammonium bicarbonate by centrifugation at 1800 x g for 3 minutes at room temperature. After removal of the supernatant, the resins were incubated in 200 μl of 50 mM ammonium bicarbonate containing 10 mM TCEP for 15 minutes at 37°C with shaking and next, the resins were alkylated with 4 μl of 0.5 M IAA (Iodoacetamide) for 15 minutes in the dark at 37°C with shaking. The samples were then digested with 1 μg of trypsin overnight at 37°C with shaking. The supernatant was transferred to a clean Eppendorf tube and acidified by adding trifluoroacetic acid to a pH of 3. The acidified samples were then desalted on Pierce^TM^ C18 stage tips (Thermo Fisher Scientific) and dried in a vacuum concentrator (SpeedVac, Eppendorf). The peptides were reconstituted in 40 μl of 0.5% formic acid and 18 μl of each sample was analysed by nanoflow liquid chromatography tandem mass spectrometry (LC-MS/MS) using an Orbitrap Elite Hybrid Mass Spectrometer (Thermo Fisher Scientific) coupled online to an UltiMate RSLCnano LC System (Dionex^TM^). The system was controlled by Xcalibur 3.0.63 (Thermo Fisher) and DCMSLink (Dionex). Peptides were desalted on-line using an Acclaim PepMap 100 C18 nano/capillary BioLC, 100A nanoViper 20 mm x 75 µm I.D. particle size 3 µm (Fisher Scientific) at a flow rate of 5 μl/min and then separated using a 125-min gradient from 5 to 35% buffer B (0.5% formic acid in 80% acetonitrile) on an EASY-Spray column, 50 cm × 50 μm ID, PepMap C18, 2 μm particles, 100 Å pore size (Fisher Scientific) at a flow rate of 0.25 μl/min. The Orbitrap Elite was operated with a cycle of one MS (in the Orbitrap) acquired at a resolution of 60,000 at m/z 400, with the top 20 most abundant multiply charged (2+ and higher) ions in a given chromatographic window subjected to MS/MS fragmentation in the linear ion trap. An FTMS target value of 1e6 and an ion trap MSn target value of 1e4 were used with the lock mass (445.120025) enabled. Maximum FTMS scan accumulation time of 100 ms and maximum ion trap MSn scan accumulation time of 50 ms were used. Dynamic exclusion was enabled with a repeat duration of 45 s with an exclusion list of 500 and an exclusion duration of 30 s.

### Mass spectrometry data analysis

Raw data collected from mass spectrometry were analysed with MaxQuant version 1.6.2.6. All data were searched against a human UniProt database. Search parameters contained digestion set to Trypsin/P with maximum of 2 missed cleavages, methionine oxidation and acetylation at N-terminal peptide as variable modifications, carbamidomethylation at cysteine as fixed modification, match between runs enabled with a match time window of 0.7 min and a 20-min alignment time window, label-free quantification (LFQ) enabled with a minimum ratio count of 2, minimum number of neighbours of 3 and an average number of neighbours of 6. A first search precursor tolerance of 20 ppm and a main search precursor tolerance of 4.5 ppm was used for FTMS scans and a 0.5 Da tolerance for ITMS scans. A protein false discovery rate (FDR) of 0.01 and a peptide FDR of 0.01 were used for identification level cut-offs.

Downstream data analysis of MaxQuant outputs was performed using Perseus version 1.5.6.0 with LFQ intensities as the main categories. LFQ intensities were transformed by log2(x) and grouped according to experimental conditions (+Tet and -Tet). LFQ Intensities were normalised by subtracting the medians and missing values were randomly imputed with a width of 0.3 and downshift of 3 from the standard deviation. To identify quantitatively changed proteins between +Tet and -Tet groups, two-tailed Student’s t-tests were performed with a S0 value of 2 and a permutation-based FDR of 0.05. Gene ontology (GO) analysis was conducted by Database for Annotation, Visualization and Integrated Discovery (DAVID) (https://david.ncifcrf.gov/) against H. sapiens proteome background. Statistical significance of enrichment of GO terms were determined by adjusted p-values < 0.05.

### Traction force microscopy

HEK293-tet-RhoA-TurboID were seeded on 35 mm 12 kPa collagen coated plates containing 0.2 µm red fluorescent beads (Cell Guidance Systems, UK) and incubated for 24 hours. 30 minutes before imaging, the media was changed to fresh media containing 0.05 µg/mL Hoechst dye. Plates were imaged using a Cell Discoverer 7 microscope (Zeiss, Germany), maintained at 37°C, 5% CO_2_. Images of cell clusters and beads were taken before and 2 hours after treatment with 1 µg/mL Tetracycline. Traction force was analysed in FIJI [71]. Bead displacement was determined using particle image velocimetry followed by Fourier transform traction cytometry to calculate traction force [72, 73].

### Preparation of polyacrylamide hydrogel for cell culture

Collagen-coated polyacrylamide (PAA) hydrogels were prepared and directly polymerized in polystyrene culture dishes as previously described [74, 75]. In brief, prior to the preparations of the PAA gel, an acrylic-based photosensitive Loctite resin (Ellsworth) was applied on the bottom of the dish to form an adhesive layer for attachment of polyacrylamide gel to culture dishes. A solution of different concentrations of acrylamide and bis-acrylamide was prepared to produce gels of various stiffness. The solution was mixed with 1% APS and 0.1% TEMED and then dispensed on Loctite resins-coated dishes. The acrylamide/bis-acrylamide solution was quickly overlayed with parafilm. After 2 hours, the parafilm was carefully removed from the polymerized PAA gels and the gels were washed with double distilled H_2_O and then three times with PBS to adjust pH and salinity within the gel. Plates used for soft conditions had a stiffness of 0.2 kPa or 0.7 kPa. Plates used for stiff conditions had a stiffness of 25kPa. Plates were custom-made as described above or commercial (Matrigen). Custom-made and commercial plates gave comparable results when tested for YAP subcellular localisation (Suppl. Fig. 3D, E).

The surface of PAA gels was functionalized by NHS-acrylate (Sigma-Aldrich) and Irgacure 2959 (Sigma-Aldrich) in PBS and exposure to UV radiation (γ=365 nm) for 15 minutes. PAA gels were washed gently once with PBS followed by coating with 100 μg/ml collagen type I (Gibco) in PBS overnight at room temperature. After washing with PBS, gels were sterilised by UV and then equilibrated three times with culture medium for 1 hour each. Cells with the amount of 2.5x10^5^ were seeded on top of the gels. Cells were cultured for 72 hours for immunofluorescence staining and for RNA or protein extraction unless otherwise stated.

### Immunofluorescence and image analysis

Immunofluorescence staining was performed by standard procedure. Briefly, cells seeded on coverslips or polyacrylamide gels were fixed with 4% paraformaldehyde in PBS for 10 minutes and then permeabilized with 0.1% Triton X-100 in PBS for 15 minutes at room temperature. After incubation in blocking buffer (0.05% FBS/0.01% Tween-20/PBS) for 1 hour at room temperature, cells were incubated with primary antibodies for 2 hours, then with Alexa Fluor conjugated secondary antibodies (Thermo Fisher Scientific) for 1 hour at room temperature. Primary antibodies were diluted in blocking buffer at the following dilution ratios: YAP 1:200 (sc-101199 Santa Cruz), YAP 1:500 (14074 Cell Signalling Technology), PTBP1 1:300 (32-4800 Thermo Fisher Scientific),) and nuclei were counterstained with Hoechst dye (Invitrogen). Both primary and secondary antibody incubation were followed by three PBS washes. To visualise immunofluorescence-stained cells on coverslips, cells were embedded in Prolong Gold anti-fade mounting medium (Invitrogen) and the images were acquired with a 60x oil immersion objective in an Olympus Fluoview 1000 Confocal microscope or an Olympus Epifluorescence microscope. To observe fluorescence images of cells on polyacrylamide gels, cells were soaked in PBS and the images were taken with a 60x water immersion objective on an Olympus Fluoview 1000 Confocal microscope. The images were processed with Fiji-ImageJ software. Fluorescence intensity of YAP and PTBP1 were assessed using Fiji software and the Nuclear/Cytoplasmic ratios of these target proteins were calculated using the following formula:

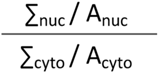

Where ∑_nuc_ and ∑_cyto_ are the sum of the intensity values of the pixels in the nucleus and cytoplasm in the images, and A_nuc_ and A_cyto_ represent the nuclear and cytoplasmic areas.

### Cell proliferation assay

5-bromo-2-deoxyuridine (BrdU) (Merck) immunofluorescence staining was performed to assess the influence of ECM stiffness on MCF10A proliferation. In brief, MCF10A cells of 50% confluency on polyacrylamide gel (Matrigen) were incubated with 20 μM BrdU (Merck) at 37°C for 3 hours. Cells were washed with PBS followed by fixation with 4% paraformaldehyde for 15 minutes. After washing, cells were incubated in a permeabilization buffer for 15 minutes followed by treatment with 2M HCl for 5 minutes and then phosphate/citric acid buffer for 20 minutes. Cells were incubated in blocking buffer (0.5%FBS, 0.01% Tween 20, 1xPBS pH 7.4) for 1 hour before being treated with anti-BrdU antibody (1:1000, Abcam) overnight at RT. After washing with PBST, cells were incubated with AlexFluor-594 conjugated secondary antibody (1:500, Thermo FIsher Scientific) and Hoechst dye (1:1000, Thermo FIsher Scientific) for 1 hour. BrdU incorporation was quantified using fluorescence microscopy (Olympus Fluoview 1000 confocal). Cell proliferation was assessed as the percentage of BrdU-positive/total number of cells.

### Reverse Transcription (RT)-PCR

The change of alternative splicing by various ECM stiffnesses was verified by RT-PCR. In brief, MCF10A were cultured on soft PAA gel for 5 days and then split and cultured either on soft or on stiff PAA gel for three days. Total RNA was isolated from MCF10A using RNeasy Mini Kit (QIAGEN) according to manufacturer’s protocols. RNA concentration and purity were measured using a NanoDrop^TM^ Lite Spectrophotometer (Thermo Fisher Scientific). After DNase I treatment, 200 ng of total RNA was reverse-tanscribed into complementary DNA with SuperScript^TM^ II Reverse Transcriptase (Invitrogen). Subsequently, the cDNA was mixed with GoTaq^®^ DNA polymerase (Promega) and primers listed in Table S3, and PCR was performed on a T100 Thermal Cycler (Bio-Rad). Annealing temperatures and extension time during the thermocycling was optimised for each primer pair and size of amplicons. PCR products were analysed by 2% agarose gel stained with SYBRE^TM^ Safe (Invitrogen) and visualised using Gel Doc EZ System (Bio-Rad). DNA bands were quantified using Fiji software. The splicing ratio of each isoform was first normalized to its DNA size and was calculated as the inclusion level (%) of the cassette exon over the sum of all isoforms.

### Statistical analysis and other software

Graphs and statistical comparison were carried out using GraphPad Prism version 9.5.1. Unpaired t-test with Welch’s correlation were used to compare two cases and ANOVA tests were performed to analyse more cases. Data were presented as mean ± s.d. Details on sample numbers and analysis methods were given in each legend. Statistical significance is denoted by asterisks*: \**p*<0.05, \*\**p*<0.01, \*\*\**p*<0.001, and \*\*\*\**p*<0.0001. Diagrams were made using Biorender and Microsoft Powerpoint.

