## Supplementary figures for "Mechanical control of the splicing factor PTBP1 regulates extracellular matrix stiffness-induced cell proliferation and mechanomemory"

### Supplemental Figures

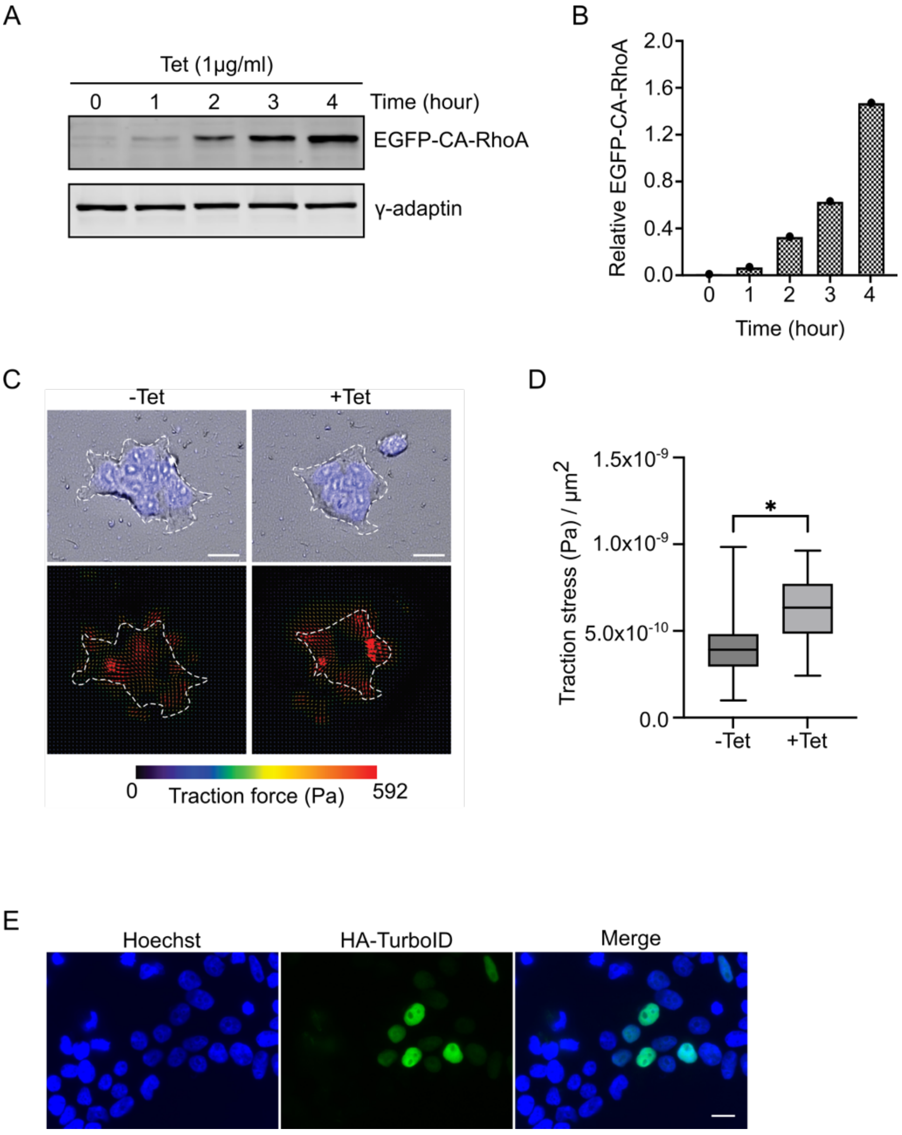


**Fig. S1 Validation of screen approach, Related to Figure 1. A:** HEK293-tet-RhoA expressed constitutively active RhoA by tetracycline induction in a time-dependent manner. γ-adaptin was used as loading control. **B:** Quantification of (A), bars represented normalised EGFP-CA-RhoA expression. **C:** Detection of traction force in HEK293-tet-RhoA before and after two hours of tetracycline treatment. Cells were visualized by staining of nuclei. Red arrows indicated the force direction. Scale bar, 20 μm. **D:** Quantification of traction stress in HEK293-tet-RhoA before and after tetracycline treatment. Data was analysed by unpaired t-test (n=3). Values are means ± s.d. *$p < 0.05$. **E:** Transient overexpression of HA conjugated NL-TurboID in HEK293-tet-RhoA showed their nuclear localization. Scale bar, 15 μm.

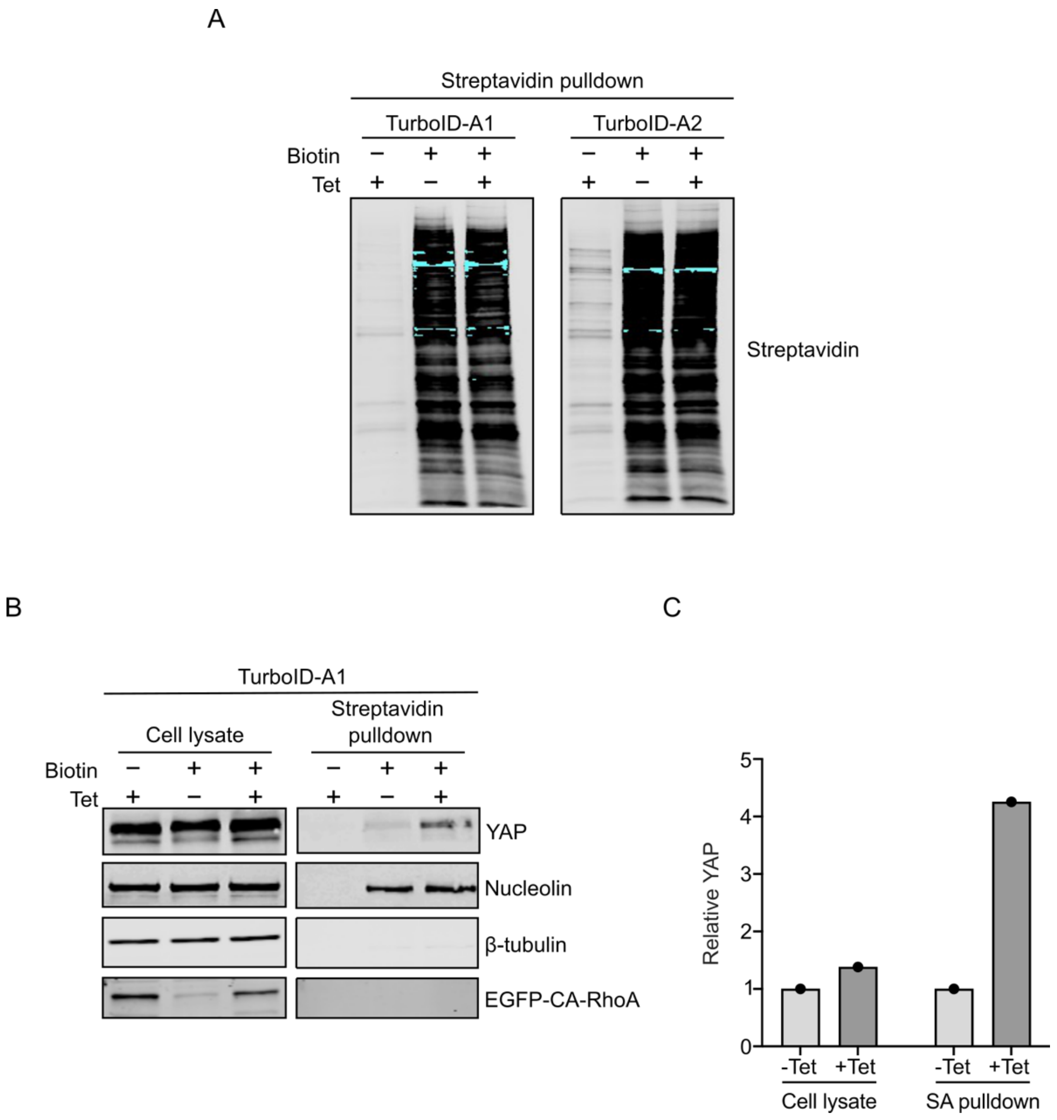


**Fig. S2 Validation of HEK-tet-RhoA-TurboID stable clones, Related to Figure 1. A:** Western blot of proteins extracted from two independent HEK-tet-RhoA-TurboID clones (TurboID-A1 and TurboID-A2) showing the biotinylated nuclear proteins in cells treated with 500 μM of biotin for 20 minutes. **B:** Western blot of indicated proteins in cell lysate and streptavidin pulldown sample from TurboID-A1 clone. Nucleolin and β-tubulin were used as nuclear and cytosolic marker respectively. **C:** Quantification of YAP expression from **B** shows increased nuclear YAP after tetracycline treatment.

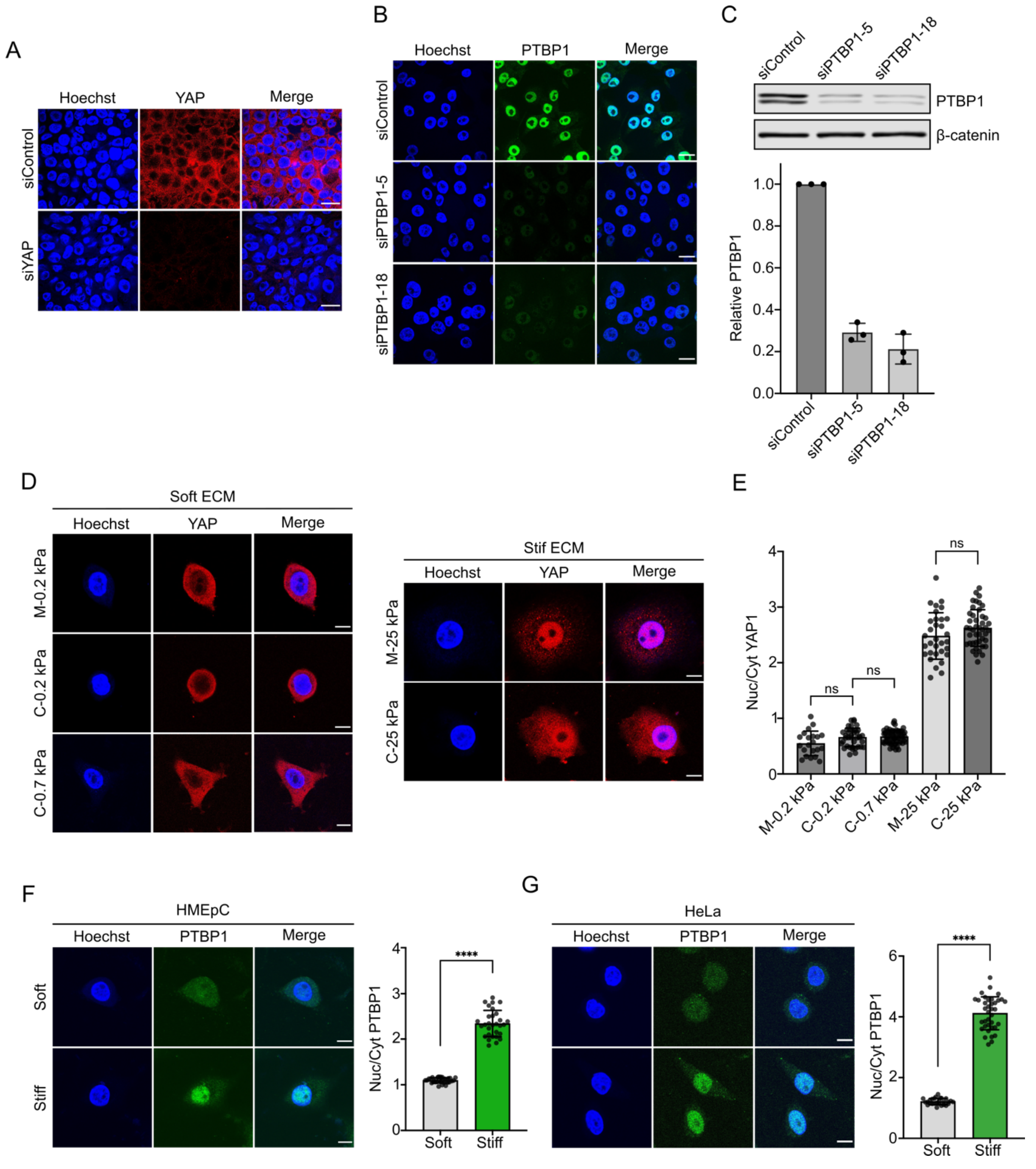


**Fig. S3 ECM stiffness regulates subcellular localization of PTBP1 and YAP, Related to Figure 3. A:** Representative immunofluorescence images of siRNA-mediated knockdown of YAP in MCF10A. **B:** Reduced PTBP1 level after siRNA-mediated knockdown in MCF10A was presented by immunofluorescence staining. **C:** Western blot confirmed knockdown efficiency carried out by indicated PTBP1 siRNA. β-catenin was used as loading control. Bar diagram shows the normalised PTBP1 level after treatment of indicated siRNA. **D:** Comparison of nuclear localization of YAP in MCF10A cultured on polyacrylamide gel of various ECM stiffness. **E:** Quantification of **D** showed the nuclear to cytosolic ratio for YAP. C: customized gel; M: Commercial Matrigen. **F-G:** Enrichment of PTBP1 in the nucleus by stiff ECM in indicated cell line. Corresponding quantification of nuclear to cytosolic ratio for PTBP1. All experiments n=3. Scale bar for **A, B**, 20 µm. Scale bar for **D, F, G**, 10 µm. Data was analysed by unpaired t-test (n=3). Values are means ± s.d. ****$p$< 0.0001. ns: not significant.

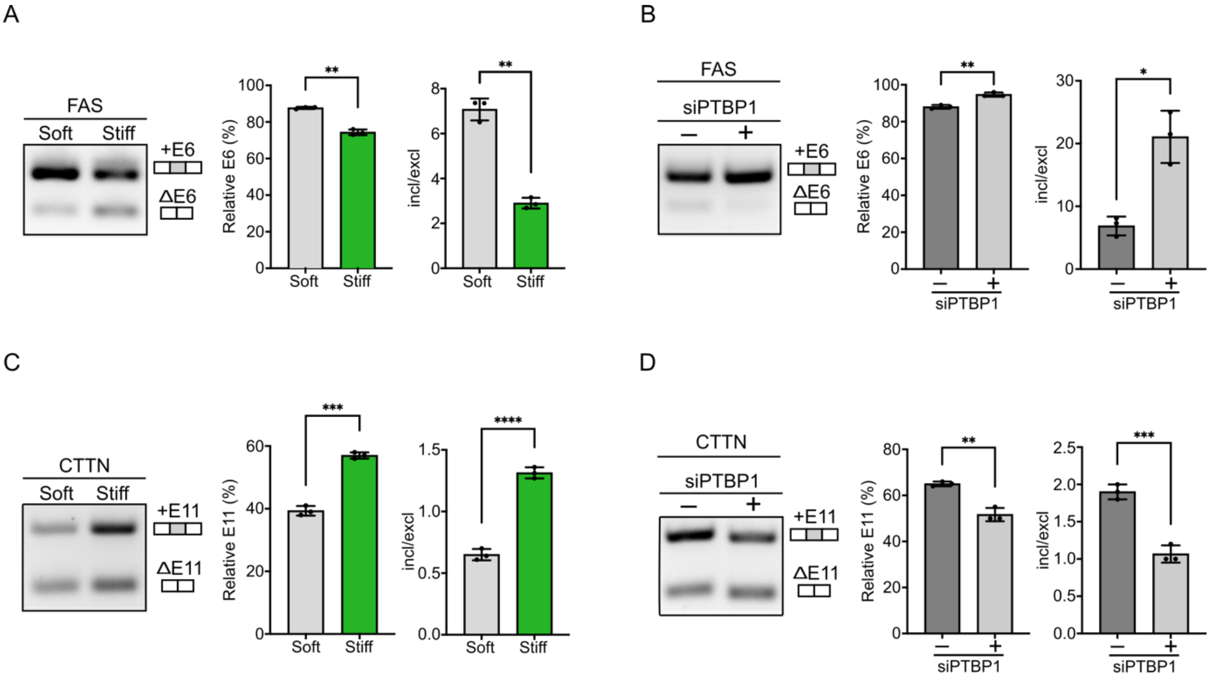


**Fig. S4 ECM stiffness modulates alternative splicing through PTBP1 activity, Related to Figure 4. A-D:** RT-PCR analysis with MCF10A showed the alternative splicing of cassette exons on FAS and CTTN regulated by ECM stiffness and PTBP1 Knockdown. Cassette exons (boxes) are indicated. Bar diagram summarises the splicing of cassette exons. The relative cassette exon (%) was defined as the ratio of exon-included isoform over the sum of exon-included and exon-excluded isoforms, while the incl/excl was calculated as the ratio of exon-included isoform against exon-excluded isoform. All experiments n=3. Statistical analyses were performed by unpaired t-test. Values are means ± s.d. *p< 0.05, **$p$< 0.01, ***$p$< 0.001, ****$p$< 0.0001.

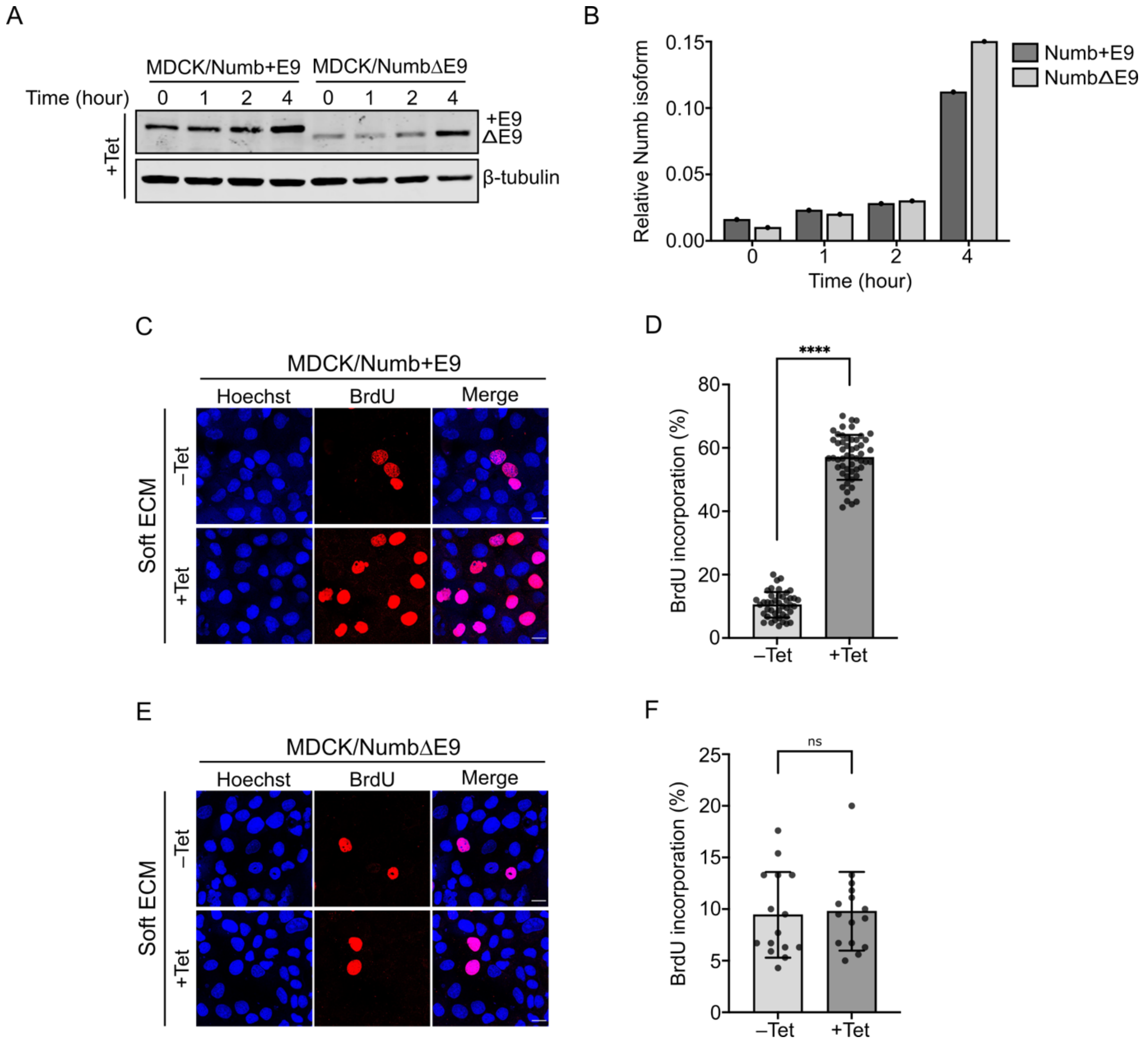


**Fig. S5 Numb E9 isoform promotes cell proliferation on soft ECM, Related to Figure 5.** **A:** Induction of Numb+E9 or NumbΔE9 isoform in two different MDCK/Numb clones by tetracycline for indicated time. **B:** Quantification of normalised Numb isoform in A. **C-D:** Cell proliferation on soft ECM was determined by BrdU assay in MDCK/Numb+E9 stable clone treated with control (-Tet) or tetracycline (+Tet). Quantification was calculated as the percentage of BrdU-positive cells (n=3). **E-F:** Cell proliferation on soft ECM was performed by BrdU assay in MDCK/NumbΔE9 stable clone without tetracycline (-Tet) or in the presence of tetracycline (+Tet). Quantification was assessed as percentage of BrdU-positive cells (n=1). Scale bar for **C**, **E**, 20 μm. Data was analysed by unpaired t-test and presented as means ± s.d. ****$p < 0.0001$. ns: not significant.
